# Exploring the role of the outer subventricular zone during cortical folding through a physics-based model

**DOI:** 10.1101/2022.09.25.509401

**Authors:** Mohammad Saeed Zarzor, Ingmar Blümcke, Silvia Budday

## Abstract

The human brain has a highly complex structure both on the microscopic and macroscopic scales. Increasing evidence has emphasized the role of mechanical forces for cortical folding – a classical hallmark of the human brain. However, the link between cellular processes at the microscale and mechanical forces at the macroscale remains insufficiently understood. Recent findings suggest that an additional proliferating zone, the outer subventricular zone (OSVZ), is decisive for the particular size and complexity of the human cortex. To better understand how the OSVZ affects cortical folding, we establish a multifield computational model that couples cell proliferation and migration at the cell scale with growth and cortical folding at the organ scale by combining an advection-diffusion model with the theory of finite growth. We validate our model based on data from histologically stained sections of the human fetal brain. Finally, we address open questions regarding the role of the OSVZ for the formation of cortical folds. The presented framework not only improves our understanding of human brain development, but could eventually help diagnose and treat neuronal disorders arising from disruptions in cellular development and associated malformations of cortical development.

## Introduction

The brain is one of the most fascinating organs in the human body. Its complex structure on both micro- and macroscopic scales closely correlates with the unique cognitive abilities of humans. Cortical folding is one of the most important features of the human brain. Still, compared to other mammals, the human brain is neither the largest nor the most folded brain. However, relative to its size, it has the largest number of cortical neurons that connect with billions of neuronal synapses ***Herculano-Houzel (2009)***. This fact attracted the attention of neuroscientists over the past few years to explore the source of these cells and how they develop in the early stages of brain development.

The number of brain cells is determined *in utero* through the proliferation process. Previous studies on different lissencephalic species such as mice have shown that cell division in the brain is confined to a small region near the cerebral ventricles ***Hansen et al. (2010)***. However, in gyrencephalic species, this seems to be slightly different. Recent findings show that the human brain, for example, is characterized by two proliferation zones with two different types of progenitor cells. Both zones produce neurons that later migrate towards the outer brain surface and form the cortex ***Lui et al. (2011)***.

In rodents, progenitor cells around the ventricular zone (VZ) generate intermediate progenitor cells as their daughters, which accumulate above the ventricular zone and form a new layer called the subventricular zone (SVZ) ***Noctor et al. (2002)***. In humans, there is an additional outer layer of the subventricular zone, often referred to as outer subventricular zone (OSVZ) ***Hansen et al. (2010)***; ***Lui et al. (2011)***; ***Noctor et al. (2007)***. This zone was first discovered in the monkey brain by Colette Dehay and his colleagues ***Smart et al. (2002)***, and confirmed in the human brain by several following studies ***Huttner and Kosodo (2005)***.The OSVZ seems to play a significant role in the proliferation process and affects the size and complexity of the human cortex. The evidence for this allegation is the wave of cortical neurogenesis that coincides with the cell division in the OSVZ ***Lukaszewicz et al. (2005)***. At the macroscopic scale, the high proliferation in the OSVZ coincides with a significant tangential expansion of the cortical layers. The latter is an essential factor for the formation of cortical folds ***Reillo et al. (2011)***. Still, it remains unknown, how exactly this proliferation process in the OSVZ affects gyrification of the forming cortex. Different approaches have been used to understand the relation between cellular mechanisms at the microscopic scale and corticogenesis at the macroscopic scale. Genetic analyses and experimental studies using cell culture models and brain organoids have given first valuable insights concerning the source of cells and their behavior ***Hansen et al. (2010)***. Here, we intend to complement these studies by using a numerical approach to bridge the scales from the behavior of different progenitor cell types at the cell scale to the emergence of cortical folds at the tissue or organ scale.

From a mechanics point of view, forces that are generated due to cellular processes may act as a link to understand the underlying mechanisms behind cortical folding ***Budday et al. (2015)***. Many previous studies tried to explain normal and abnormal cortical folding either from a purely biological or mechanical perspective. However, it will not be possible to capture the folding mechanism without considering both perspectives at the same time ***Zarzor et al. (2021)***. In other words, to fully understand the physiological and pathological mechanisms underlying cortical folding in the developing human brain, we need to study the coupling between cellular processes and mechanical forces – to eventually assess how disruption of cellular processes affect the folding pattern and lead to malformations of cortical development ***Guerrini et al. (2008)***; ***Blumcke et al. (2021)***; ***Llinares-Benadero and Borrell (2019)***.

To fill this knowledge gap, we establish a two-field computational model that accounts for both proliferating zones in the human brain. The first field in the model describes the cellular processes occurring during human brain development, where we use an advection-diffusion equation to mimic the migration in the subcortex and neuronal connectivity in the cortex ***de Rooij and Kuhl (2018)***; ***Zarzor et al. (2021)***. We add two source terms to consider the division in both zones, VZ and OSVZ. Regarding the second field, we use the theory of finite growth. Finally, we validate the model through a comparison of the simulation results with histologically stained sections of the human fetal brain (HBS) with regard to the cell density distributions and surface morphology.

## Cellular processes during brain development

The central cellular unit that plays a critical role in essential processes of brain development is a type of progenitor cells called radial glial cells (RGCs). In the early stage of brain development, when neurogenesis begins, the neuroepithelial cells transform into RGCs ***Noctor et al. (2007)***. Around gestational week (GW) five, these cells locate near cerebral ventricles, in the ventricular zone (VZ), where they undergo interkinetic nuclear migration (INM). The associated symmetric division behavior leads to a significant increase in the number of RGCs and results in both increased thickness and surface area of the VZ ***Blows (2003)***; ***Fish et al. (2008)***; ***Bystron et al. (2008)***. Subsequently, the cells switch to an asymmetric division behavior and generate intermediate progenitor cells (IPCs) ***Noctor et al. (2004)***.The IPCs migrate to the subventricular zone (SVZ) where they proliferate and produce neurons, as illustrated in Figure 1 ***Noctor et al. (2007)***; ***Pebworth et al. (2021)***. Ultimately, the majority of cortical neurons are produced by IPCs ***Lui et al. (2011)***; ***Libé-Philippot and Vanderhaeghen (2021)***.

**Figure 1.**
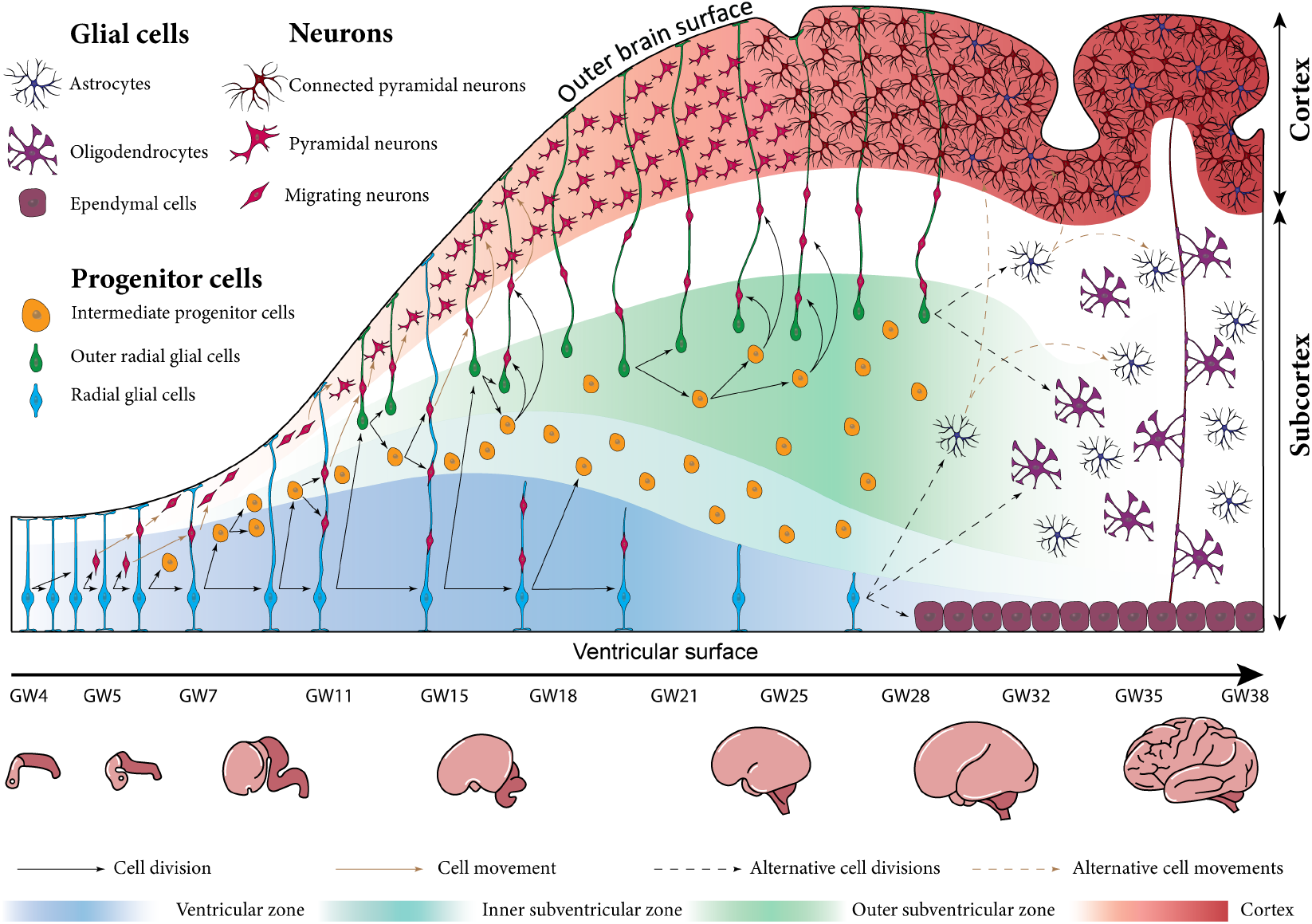
Schematic illustration of human brain development between gestational weeks 4 and 38 at the cellular scale (top) and the organ scale (bottom). In the early stage of development, the repetitive division of radial glial cells (RGCs) in the ventricular zone (VZ) significantly increases the total number brain cells. The newly born intermediate progenitor cells (IPCs) accumulate above the VZ and form a new layer called the inner subventricular zone (ISVZ). The outer radial glial cells (ORGCs) that are produced around gestational week (GW) 11 form a new layer called the outer subventricular zone (OSVZ). The neurons generated from progenitor cells migrate along RGC fibers towards the cortex. Around GW 28, the migration process is almost finished, and the RGCs switch to produce different types of glial cells like astrocytes and oligodendrocytes.

According to the radial unit hypothesis proposed by Pasko Rakic over thirty years ago, the radial glial cell fibers organize the migration process, which starts around GW six ***Nonaka-Kinoshita et al. (2013)***. He postulated that these fibers form a scaffold to guide neurons during their migration from the proliferating zones to their final destination in the cortex, which forms the outer brain surface ***Rakic (1988)***; ***Luietal.(2011)***. In the gyrencephalic species, those fibers have a characteristic fan-like distribution ***Borrell and Götz (2014)***; ***Nonaka-Kinoshita et al. (2013)***. However, it is still under debate whether this unique distribution is the cause or rather the result of cortical folding. Our recent computational analyses support the latter, i.e., that it is the result of cortical folding ***Zarzor et al. (2021)***. The migration process synchronizes with a radial expansion of all brain layers. Still, the VZ does not expand remarkably as the IPCs move out to the SVZ. The migrated neurons organize themselves in the six-layered cortex in an inside-out sequence, where the early-born neurons occupy the inner layers ***Gilmore and Herrup (1997)***. Until GW 23, the outer brain surface is still smooth, although the bottom four layers of the cortex are already filled with neurons ***Shinmyo et al. (2017)***. The first brain folds begin to form around GW 25 as the cortical layer significantly expands tangentially ***Budday et al. (2015)***. Importantly, at GW 25, the cortical neuronal connectivity emerges and comes along with the horizontal elongation of neuronal dendrites ***Takahashi et al. (2012)***.

While the processes summarized above are common among mammals, the human brain has some specific features that play a significant role in increasing the number of cortical neurons, and enhancing the complexity of cortical folds ***Libé-Philippot and Vanderhaeghen (2021)***. At the beginning of the second trimester, around GW 11, the original RGCs switch from producing the IPCs to producing a special kind of cells that are found in all gyrencephalic species but are enriched in the human brain. The newly generated cells are similar to the RGCs in terms of shape and function, but unlike the original RGCs, they migrate to the outer layer of the subventricular zone (OSVZ) after they are born. Therefore, they are called outer radial glial-like cells (ORGCs) ***Lui et al. (2011)***; ***Fietz et al. (2010)***; ***Hansen et al. (2010)***; ***Reillo et al. (2011)***; ***Nonaka-Kinoshita et al. (2013)***. We would like to note that some literature refers to this type of cells as basal radial glial cells. While the original RGCs have a bipolar morphology with two processes – one extending to the cerebral ventricle and one to the outer cortical surface –the ORGCs have a distinct unipolar structure with only a single process extending to the outer cortical surface ***Hansen et al. (2010)***; ***Betizeau et al. (2013)***; ***Reillo et al. (2011)***; ***Nonaka-Kinoshita et al. (2013)***.

The OSVZ shows a significantly more pronounced radial expansion compared to the inner sub-ventricular zone (ISVZ) and VZ between GWs 11.5 and 32. The immediate reason causing this difference is the characteristic division behavior of ORGCs: they translocate rapidly in radial direction before they divide, which scientists refer to as “mitotic small translocation (MST)”***Fietz et al. (2010)***. Importantly, the MST behavior pushes the boundary of the OSVZ outward, which increases the capacity of the OSVZ to produce new neurons. The IPCs have enough space to undergo multiple rounds of division before producing neurons, which increases the overall number of generated neurons ***Kriegstein et al. (2006)***; ***Lui et al. (2011)***.

The ORGCs, like RGCs, play an important role in the proliferation process: they divide symmetrically and asymmetrically to produce further ORGCs and IPCs ***Libé-Philippot and Vanderhaeghen (2021)***. IPCs divide to generate a pair of neurons ***Lui et al. (2011)***. According to previous studies, 40% of produced neurons are generated by ORGCs at GW 13, but this ratio increases to 60% by GW 14 and exceeds 75% by GW 15.5, then after GW 17, the ORGCs become the only source of cortical neurons in the upper cortical layers ***Hansen et al. (2010)***. Beside their role in increasing the number of neurons, the ORGCs generate additional scaffolds that elongate to the outer brain surface and serve as paths for neuronal migration ***Llinares-Benadero and Borrell (2019)***; ***Nonaka-Kinoshita et al. (2013)***. Compared to other mammals, the neurogenesis of the human cortex thus divides into two main stages. The first stage is characterized by migration along a continuous scaffold consisting of the RGC fibers, which run from the ventricular surface to the outer cortical layer around GW 15. During the second stage, the migration path switches to a discontinuous form. After GW 17, the RGC fibers run from the ventricular surface to the ISVZ, while ORGC fibers run from the OSVZ to the outer brain surface, as demonstrate in Figure 1 ***Nowakowski et al. (2016)***. Consequently, the migrating neuron follows a sinuous path through numerous radial fibers before it reaches its final location in the cortex ***Lui et al. (2011)***. The newly generated scaffolds are not only important for neuronal migration, but also for the tangential expansion of the cortex. Previous studies show that a reduced number of ORGCs leads to reduced tangential expansion. In these cases, the cortex is less folded or even lissencephalic ***Poluch and Juliano (2015)***. In contrast, increasing the number of ORGCs leads to more excessive folding ***Florio et al. (2017)***; ***Borrell (2018)***. However, it is still unknown whether these effects are a result of the specific proliferation behavior of ORGCs or the associated scaffold of ORGC fibers. What is known, though, is that the existence of ORGCs is a necessary but not sufficient condition for cortical folding ***Llinares-Benadero and Borrell (2019)***.

Around GW 28, the migrating neurons occupy the first (top) cortical layer while the migration, and proliferation processes come to an end. After finishing their role during the neurogenesis stage, RGCs and ORGCs switch to produce different types of glial cells, e.g., for astrocytes and oligodendrocytes ***Schmechel and Rakic (1979)***. Also, RGCs may convert later to ependymal cells that locate around the cerebral ventricles. The oligodendrocytes form myelin sheaths around neuronal axons, wherefore the subcortical layer gains its characteristic white color.

## Computational model

To numerically study the effect ofthe ventricular zone (VZ) and the outer subventricular zone (OSVZ) on the resulting folding pattern, we simulate human brain development by using the finite element method. The influence of various factors on the emergence of cortical folds can be best shown on a simple two-dimensional quarter-circular geometry ***Darayi (2021)***, as illustrated in Figure 2A. In the following, we introduce the main equations describing the coupling between cellular mechanisms in different proliferating zones and cortical folding, which we solve numerically.

**Figure 2.**
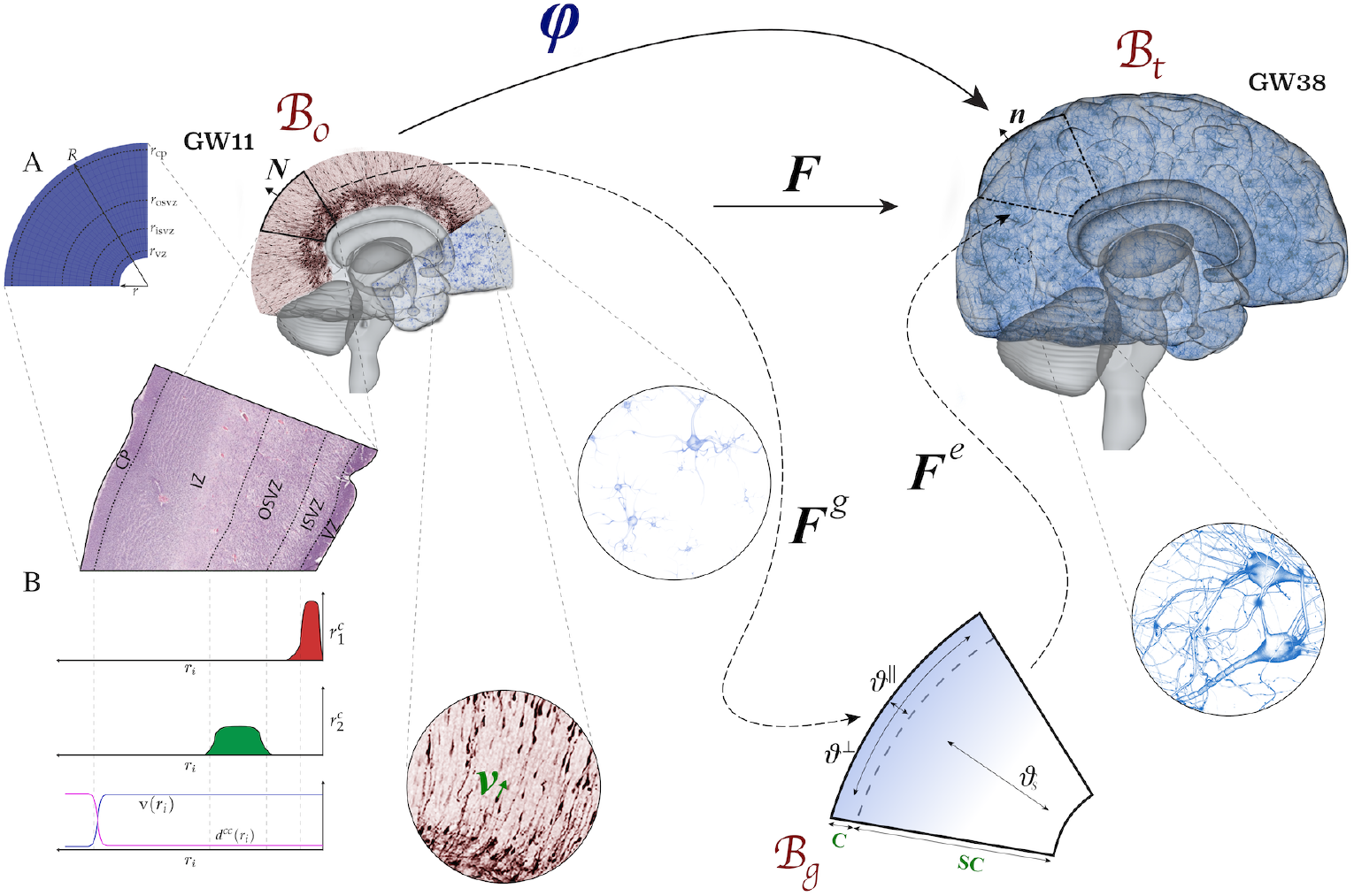
Kinematics of the multifield brain growth model. The reference configuration 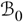 represents the initial state of the brain at gestational week (GW) 11. The spatial configuration 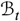 represents the state of the brain at any time *t* during development. The stress-free (intermediate) growth configuration 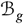 is inserted between reference and spatial configurations. (A) Simulation domain representing a part of the human brain’s frontal lobe. (B) Distribution of model parameters (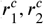, v, and *d^cc^*) along the brain’s radial direction *r_i_* from the ventricular surface to the outer cortical surface.

### Kinematics

To mathematically describe brain growth, we use the theory of nonlinear continuum mechanics supplemented by the theory of finite growth. The initial state of the brain at an early stage of development, around gestational week (GW) 11 is represented by the reference configuration 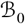. The state ofthe brain at time ***t*** later during development is represented by the spatial configuration 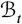. The deformation map *x* = *φ*(*X*,*t*) maps a reference point 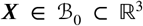 to its new position 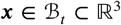 at a specific time *t*, as illustrate in Figure 2. The derivative ofthe deformation map with respect to reference point position vector is called deformation gradient ***F*** = ∇_x_*φ*. The local volume change of a volume element is described by the Jacobian ***J*** = det ***F***.

Following the theory of finite growth, we introduce a stress-free configuration between reference and spatial configuration, the growth configuration 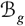. Accordingly, the deformation gradient is multiplicatively decomposed into an elastic deformation tensor ***F***^*e*^ and a growth tensor ***F***^*g*^, such that,

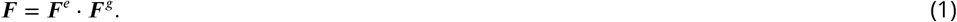

The elastic deformation tensor describes the purely elastic deformation of the brain under the effect of external forces or forces generated internally to preserve tissue continuity. On the other hand, the growth tensor controls the amount and directions of unconstrained expansion. We note that the elastic deformation tensor is reversible, while the growth tensor is not.

Besides the deformation map, we introduce the spatial cell density *c*(***x**,t*), which is a scalar independent field that depends on the spatial point position and time. It represents the number of cells per unit area ***de Rooij and Kuhl (2018)***.

For the two unknown fields, the deformation and the cell density, we introduce appropriate balance and constitutive equations in the following that then allow us to predict their evolution in space and time through numerical simulations. In the following sections, we explain how we mathematically describe the cellular processes and the mechanical problem. Then, we introduce how those are linked through the growth problem.

### Cell density problem

We formulate the balance equation of the cell density problem in such a way that we can mathematically describe the different cellular processes occurring at the microscopic scale. Temporal changes in the cell density field are kept in balance by source and flux terms. The balance equation given in the spatial configuration 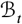 follows as

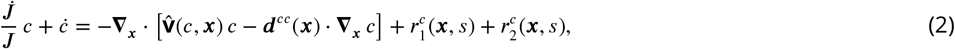

where the first flux term 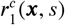 represents the migration in the subcortical plate, the second flux term ***d**^cc^* (***x***) · **∇*_x_c*** represents the neuronal connectivity in the cortex, the first source term 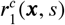 represents cell proliferation in the VZ, and the second source term 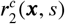 cell proliferation in the OSVZ. The migration velocity vector 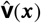 guides the cells along radial glial cell (RGC) fibers and controls their speed,

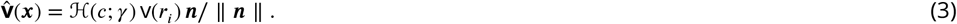

The nonlinear regularized Heaviside function 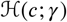 links the migration speed with the cell density field, where 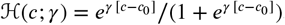. Accordingly, the cells start to migrate only when their density exceeds the critical threshold *c*_0_. The value v specifies the maximum migration speed of each individual cell in the domain. To ensure that this value vanishes smoothly at the cortex boundary ***r**_cp_*, we formulate it as a function of the radial position *r_i_*, as shown in Figure 2B, such that

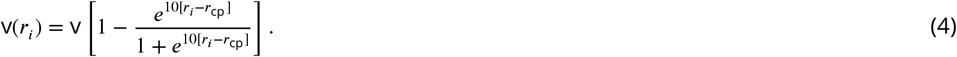

The migration direction for each cell is determined by the norm vector ***n*** that denotes the normalized RGC fiber direction in the spatial configuration. After the cells reach the cortex, they diffuse isotropically, where the diffusion tensor *d^cc^*(*x*) = *d^cc^*(*r_i_*)*I* with the diffusivity *d^cc^* and the second order unite tensor ***I*** organizes this process. In addition, we introduce the diffusivity as a function of the radial position *r_i_*, to act only in the cortex,

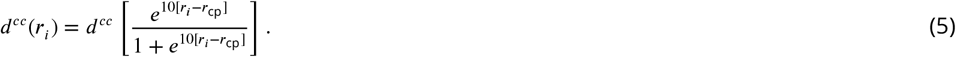

The first source term 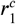 represents the RGC proliferation in the VZ, as demonstrated in Figure 2B, and is given as

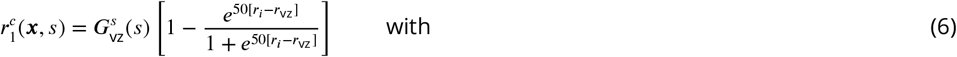

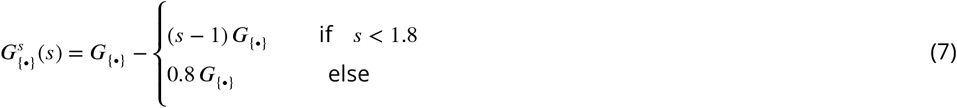

where *r*_vz_ is the outer radial boundary of the VZ. By applying equation 7 for the VZ, we ensure that the division rate decreases from its intial value *G*_vz_ to a smaller value with increasing maximum stretch value *s* in the domain, i.e., with increasing gestational age. Besides the proliferation of RGCs around cerebral ventricles in the VZ, the outer radial glial cells (ORGCs) proliferate in the OSVZ. To capture this effect, we add a second source term 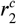, as demonstrated in Figure 2B. The second source term is given as

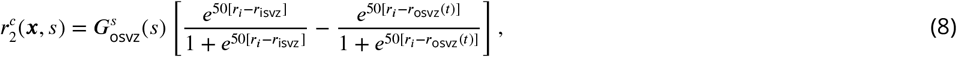

where *r*_isvz_ is the outer radial boundary of the inner subventricular zone (ISVZ). To numerically capture the expansion of the OSVZ under the effect of mitotic small translocations (MST) of ORGCs, we formulate the outer radial boundary of the OSVZ as a function to time, such that, *r*_osvz_ = *r*_isvz_ + *m*_mst_ t, where *m*_mst_ is introduced as the MST factor. Again, we apply equation 7 for the OSVZ, but in this case with the initial division rate *G*_osvz_.

### Mechanical problem

To govern the mechanical problem, we use the balance of linear momentum given in the spatial configuration 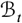

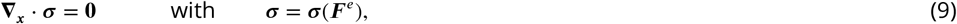

where ∇_*x*_ is the spatial gradient operator and *σ* is the Cauchy stress tensor formulated in terms of the elastic deformation tensor. The Cauchy stress is computed by deriving the strain energy function *ψ_g_* with respect to elastic deformation tensor,

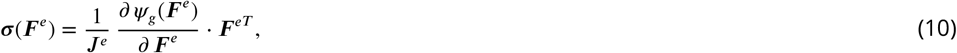

where ***J**^e^* = det ***F**^e^*. The strain energy function describes the material behavior of brain tissue mathematically. In our case, we consider a nonlinear hyperelastic material model as viscous effects, which have been observed for higher strain rates, become less relevant in the case of the slow process of brain development occurring over the course of weeks and months. Our previous studies have shown that the isotropic neo-Hookean constitutive best represents the material behavior of brain tissue during cortical folding ***Budday et al.(2020)***. The corresponding strain energy function *ψ_g_* is given as

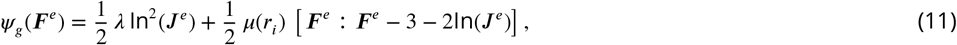

where *μ* and *λ* are the Lamé parameters. As there is a smooth transition from the cortex to the subcortical plate with distinct mechanical parameters, we use the following function

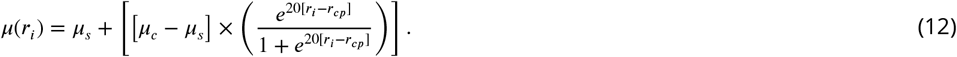

Our recent simulation study suggested that the cortical stiffness continuously changes during human brain development due to the changes in the local microstructure ***Zarzor et al. (2021)***. Accordingly, we formulate the cortical shear modulus *μ_c_* as a function of the cell density,

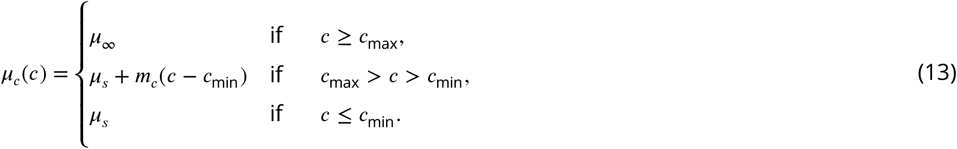

It varies in the range *μ_c_*(*c*) ∈ [*μ_s_*, *μ*_∞_], while the subcortical shear modulus *μ_s_* remains constant. The slope is defined as *m_c_* = *μ*_∞_ – *μ_s_*/*c_max_* – *c_min_* and the stiffness ratio as *β_μ_* = *μ*_∞_/*μ_s_*.

### Mechanical growth problem

The growth tensor introduced in the **Kinematics** section is the key feature in our model that links the cell density problem with the mechanical problem. As it controls the amount and direction of growth, we need to consider how cellular processes affect the physiological growth behavior in order to find an appropriate formulation. During cellular migration, the subcortical layers expand isotropically. Then, under the effect of neuronal connectivity, the cortex grows – more pronounced in circumferential than in radial direction – as illustrated in Figure 2. Thus, we introduce the growth tensor as

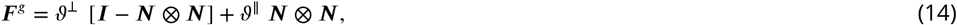

where ***N*** is the norm vector in the reference configuration 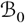 (it is linked to the spatial norm vector through ***N*** = *F*^-1^ *n*), while *ϑ*^⊥^ and *ϑ*^||^ denote the growth multipliers in circumferential and radial direction, respectively. Those multipliers control the amount of growth as a function of the cell density,

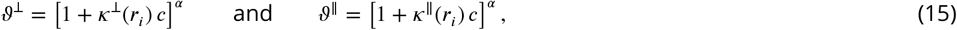

where *κ^⊥^* and *κ*^||^ are the growth factors in the circumferential and radial direction, respectively, and *α* is the growth exponent. To ensure isotropic growth in the subcortical layers, we formulate those factors as a function of the radius *r_i_*, such that

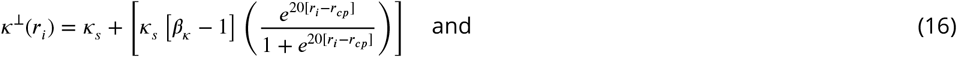

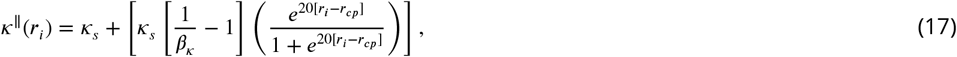

where *κ_s_* is the growth factor in the subcortical layers, and *β_κ_* is the growth ratio between *κ*^⊥^ and *κ_s_* ***Zarzor et al. (2021)***.

## Model parameters

In this work, we will consider two different cases regarding the mechanical model: The first case considers a varying cortical stiffness (VS) as introduced in the **Mechanical elastic problem** section, while the second case assumes a constant cortical stiffness (CS), i.e., *μ_c_* = *μ*_∞_ = constant. While our previous study had suggested that the simulations with varying cortical stiffness lead to morphologies that better agree with those in the actual human brain ***Zarzor et al. (2021)***, we still consider both cases in the following, VS and CS, as the situation might change when including the OSVZ and we aim to investigate corresponding interdependency effects. Table 1 summarizes the model parameters that are used in the simulation. We have previously thoroughly studied the effect of the stiffness ratio on the resulting folding pattern ***Zarzor et al. (2021)***. Here, we choose a stiffness ratio of 8 for the constant stiffness case and a ratio of 3 for the varying stiffness case. Those values led to the best agreement of simulation results with data from stained histological sections regarding the local gyrification index (LGI) value and the thickness ratio between gyri and sulci. For more details, we refer to ***Zarzor et al. (2021)***. We note that the tissue shows a stiffer behavior in the case of constant stiffness than in the case of varying stiffness for the same value of the stiffness ratio. For that reason, a higher stiffness ratio (lower stiffness in the subcortical layers since the final cortical stiffness *μ*_∞_ is constant in both cases) is required in the case of constant stiffness to achieve a similar level of folding.

**Table 1.**
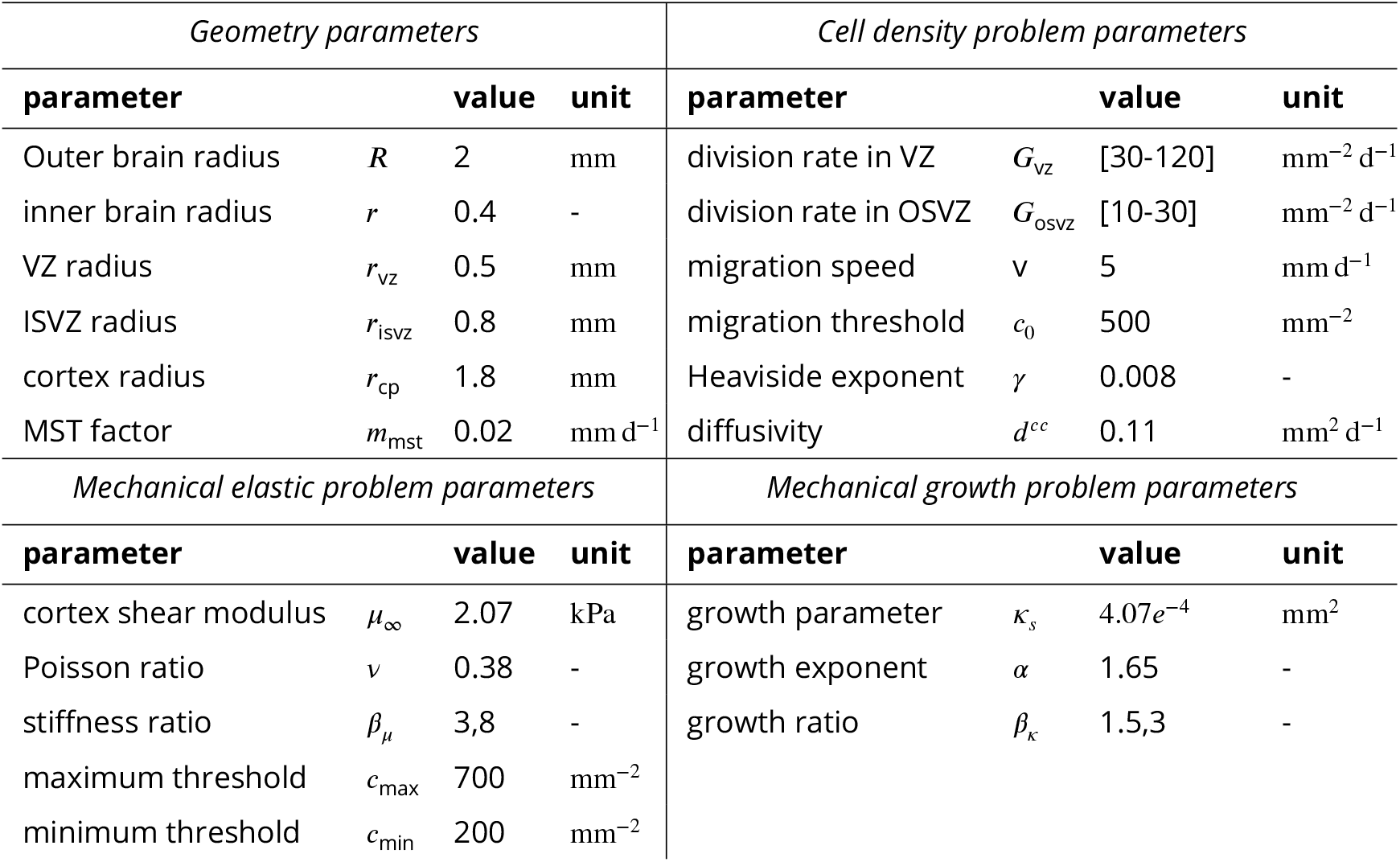
Model parameters

### Model validation

We validate our computational model by comparing the simulation results with histologically stained sections of the human fetal brain (HBS). For details on the corresponding preparation and staining, we refer to ***Zarzor et al. (2021)***. The sections belong to human fetuses aborted at gestational weeks (GW) 17, 24, 30, and 34. After defining the areas that are representative of the simulation domain introduced in the previous section (see Figure 2A) for each GW, we detect and assess the cell density using the software *Qupath*. This study was approved by the ethics review board of the University of Erlangen, and all procedures were conducted in accordance with the Declaration of Helsinki. In the following, we summarize the steps we followed to determine the cell density in human fetal brain sections (HBS). Since this analysis is intended to mainly serve as a means of comparison between different gestational weeks, a more detailed analysis using individual markers was not necessary at this stage.

1. Identify regions of interest to which the analysis should be applied, as shown in Figure 3A.
2. Pre-process the annotated area to ensure better cell detection through image color transfer, contrast ratio modification, and stain vector settings.
3. Automatically detect the cells in the relevant area by using the “positive cell detection” command in *Qupath*, which distinguishes between cell types according to the staining. Here, the negative cells (blue) are mostly glial cells or extracellular matrix components, while positive cells (red) are mostly neurons or progenitor cells, as shown in Figure 3B. While this distinction between cell types might not be fully accurate, the results are satisfactory for our specific application.
4. Count the nearby detections in a small circle with an arbitrarily chosen radius ξ = 100 μm around each detected positive cell (i) to determine the corresponding cell density by dividing the number of nearby detections by the circular area, as demonstrated in Figure 3C.
5. Visualize the cell density, as illustrated in Figure 3A.

**Figure 3.**
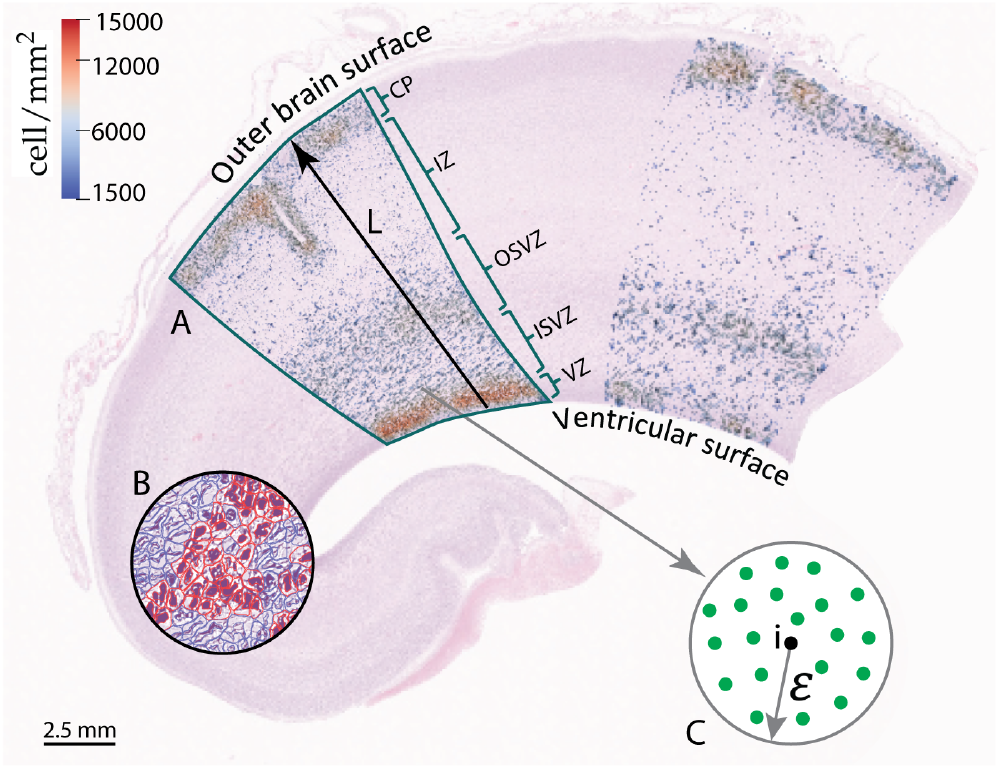
Part of the frontal lobe of a histologically stained section of the human fetal brain at gestational week (GW) 17. (A) Annotated area with final cell density distribution. (B) Example of cell detection by using *Qupath:* red cells depict neurons and blue cells glial cells. (C) Procedure to determine the cell density.

Figure 3 shows the cell density distribution around GW 17 and demonstrates the densely packed ventricular zone (VZ) due to the high proliferation rate of radial glial cells (RGCs), with about 13 600 cells/mm^2^. In addition, we can locate the other zones introduced in Figure 1. The outer subventricular zone (OSVZ) shows a higher cell density (approximately 9550cell/mm^2^) than both the inner subventricular (ISVZ) and the intermediate zone (IZ). This zone is composed of several types of cells, the original intermediate progenitor cells (IPCs) that migrated from the VZ, migrated neurons, ORGCs that produce more IPCs, and newborn neurons that are produced through the IPCs’ asymmetric division. Our analysis of the HBS (see Figure 3) also shows that the OSVZ is 1.5 times thicker than the ISVZ, even though it did not emerge at an earlier stage of development (the ISVZ emerges around GW 7 and the OSVZ around GW 11). This implies that the thickness of the OSVZ increases with time. The IZ is characterized by a low cell density with about 1600 cells/mm^2^. Still, it is a transit area for the migrating neurons. The higher cell density in the ISVZ 4800 cells/mm^2^ corroborates the presence of another type of cell besides migrating neurons, i.e. IPCs. The migration process in GW 17 is still ongoing – the cortex is not yet fully developed and filled with neurons with only about 12000cells/mm^2^. Therefore, at this stage of development, the VZ still has the highest cell density in the brain.

Figure 4 compares the cell density distribution along a line from the ventricular surface to the outer brain surface in the human fetal brain at GW 17 (indicated by line L in Figure 3) with different simulation results. For better comparability and to avoid differences in the dimensions between the HBS and simulation domain, we normalize the domain’s radius according to the extension from the ventricular to the outer brain surface in the HBS. In addition, we normalize the cell density with respect to their maximum value in the cortex. According to our previous work ***Zarzor et al. (2021)***, we include a varying stiffness in the cortex during human brain development in our simulations, adopt a stiffness ratio of 3, and a division rate in the VZ of 120.

**Figure 4.**
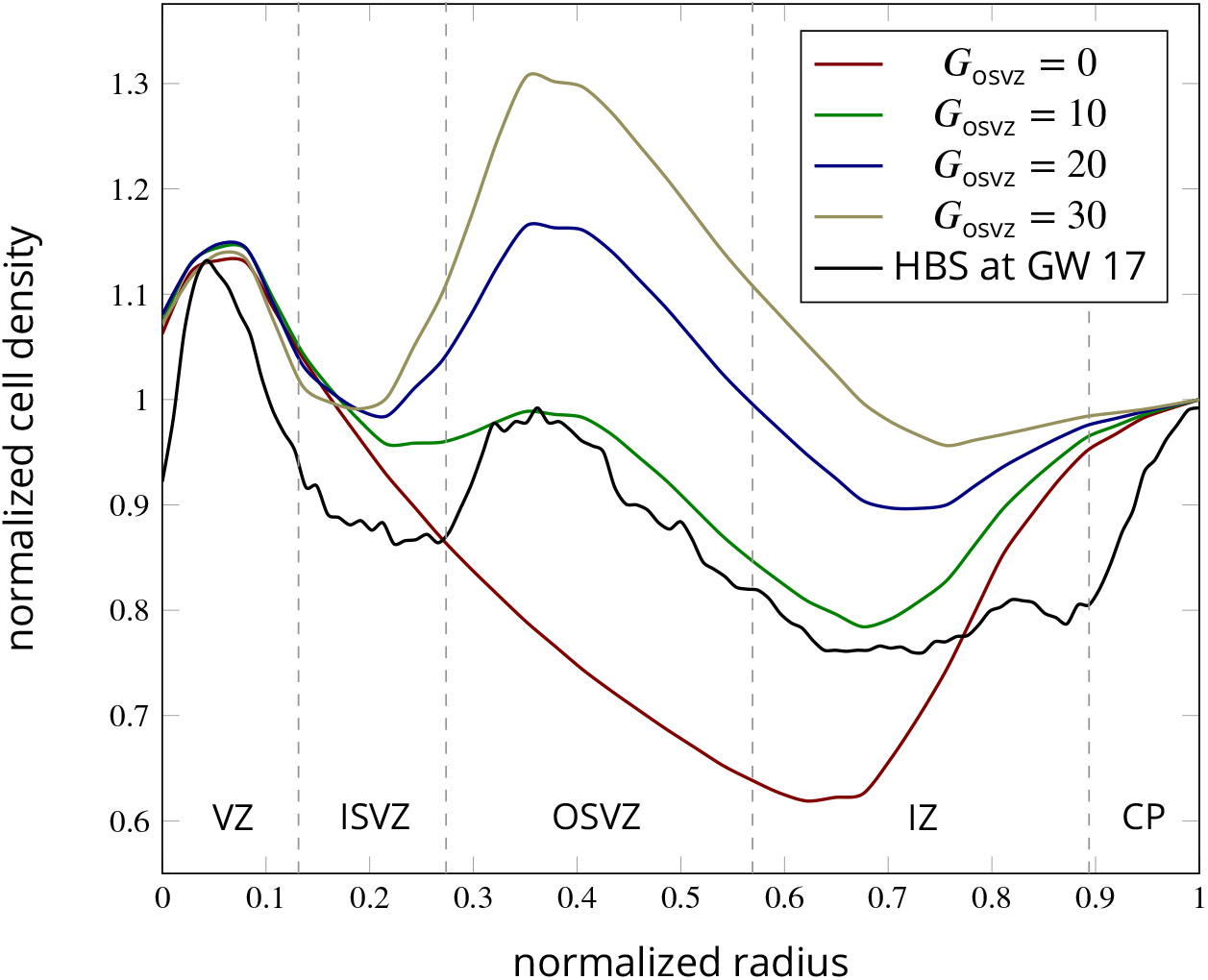
Evolution of the normalized cell density in the normalized radial direction from the ventricular surface to the outer cortical surface for numerical simulations and histologically stained human brain sections (HBS) at gestational week (GW) 17. The simulation results correspond to the time-varying cortical stiffness case with a stiffness ratio of 3, and a ventricular zone division rate of 120. The HBS results correspond to line L in Figure 3.

The simulation results for an initial division rate in the OSVZ of *G*_osvz_ = 10 well capture the trends observed in the HBS. The cell density shows a first local peak representing the VZ for a normalized radius of approximately 0.05, which is an accurate prediction. It then gradually decreases to reach its first local minimum in the ISVZ for a normalized radius of 0.3. This effect is less pronounced in the simulations than in the actual human brain. The curves start to rise again in the OSVZ to reach the second peak for a normalized radius of 0.4. Again, the simulation results capture this peak quite accurately. The second local minimum represents the IZ for a normalized radius of 0.7, while the third peak represents the cortex. The simulation results for *G*_osvz_ = 20 and 30 result in a higher cell density in the OSVZ than in the actual fetal human brain. In contrast, the curve for *G*_osvz_ = 0 (i.e., without including the effect ofthe OSVZ) shows a significantly decreased cell density in the OSVZ and IZ

## Results and discussion

In this section, we apply our computational model to answer some ofthe major questions regarding the role of the outer subventricular zone (OSVZ) for cortical folding during human brain development. The model parameters used here are introduced in the **Model parameters** section. As we mentioned above, we consider both cases constant (CS) and varying cortical stiffness (VS).

### How does cell proliferation in the OSVZ affect cortical folding patterns?

Many previous studies have tried to address this point ***Hansen et al. (2010)***. However, the experimental approach does not give a real answer to this question. In our model, we can apply different values ofthe division rate in OSVZ (*G*_osvz_) to show how the ORGCs proliferation reflects on cortical folding. Figure 5 shows the folding patterns emerging at gestational week (GW) 36 for both varying (VS) and constant (CS) stiffness cases and different values of the initial division rate in the OSVZ *G*_osvz_. In the case of CS, the sulcus becomes deeper with increasing division rate. In the case of VS, there is a more noticeable change in the folding patterns depending on the division rate *G*_osvz_. In general, the distance between neighboring sulci decreases with increasing *G*_osvz_. In addition, we observe period-doubling patterns emerge, which are most pronounced at a value of *G*_osvz_ = 20. This indicates that the proliferation in the OSVZ enhances secondary mechanical instabilities and leads to more complex folding patterns. Besides the direct relation between the proliferation in the OSVZ and the folding morphology, there are indirect effects and other aspects concerning outer radial glial cell (ORGC) proliferation, which will be discussed in more detail in the following sections. In the case of CS, the sulcus becomes deeper with increasing division rate. In the case of VS, there is a more noticeable change in the folding patterns depending on the division rate *G*_osvz_. In general, the distance between neighboring sulci decreases with increasing *G*_osvz_. In addition, we observe period-doubling patterns emerge, which are most pronounced at a value of *G*_osvz_ = 20. This indicates that the proliferation in the OSVZ enhances secondary mechanical instabilities and leads to more complex folding patterns. Besides the direct relation between the proliferation in the OSVZ and the folding morphology, there are indirect effects and other aspects concerning outer radial glial cell (ORGC) proliferation, which will be discussed in more detail in the following section.

**Figure 5.**
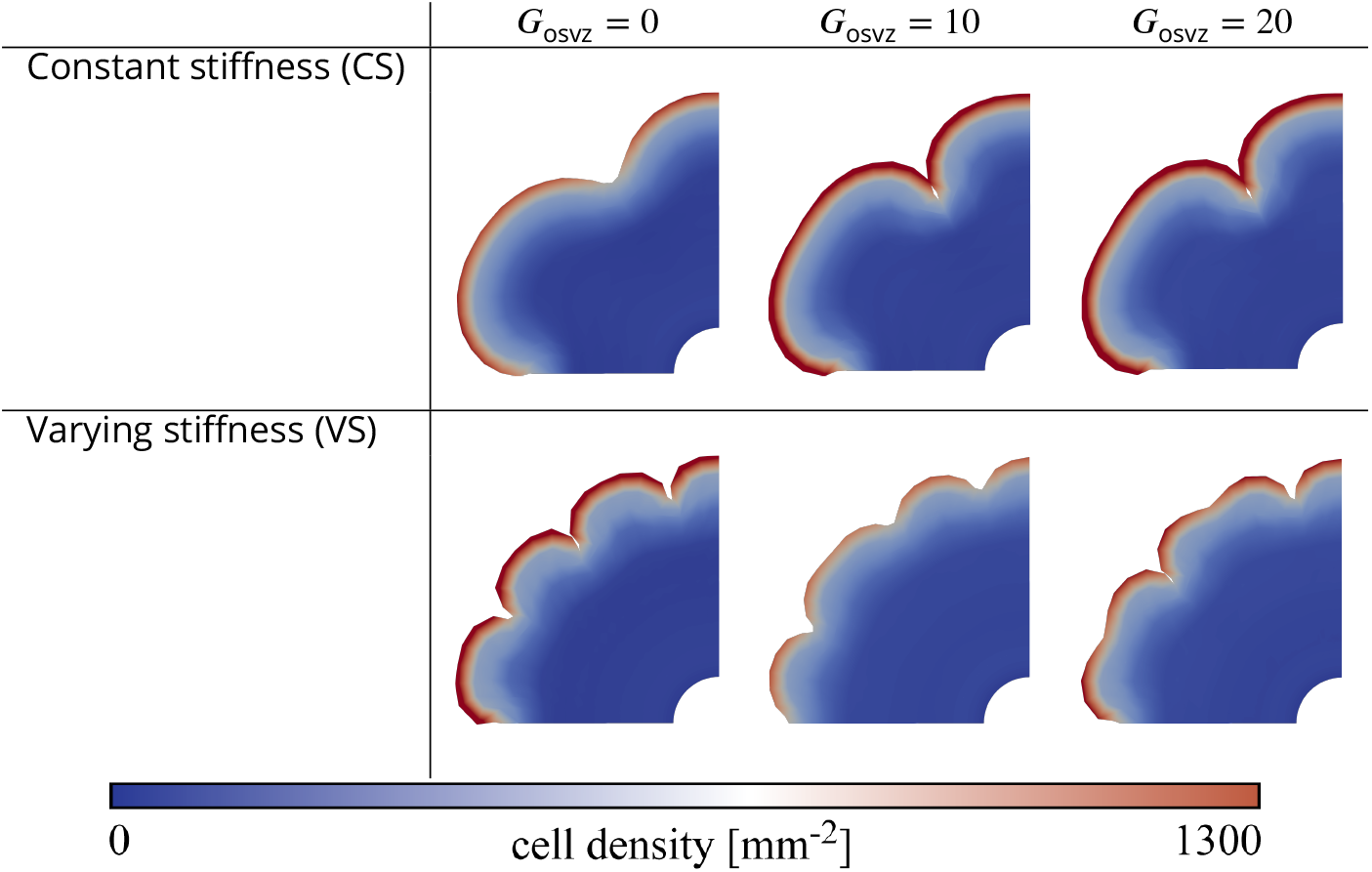
Final folding patterns at gestational week (GW) 36 for different values of the division rate in the outer subventricular zone (OSVZ) *G*_osvz_ for the constant (CS, top) and time-varying (VS, bottom) cortical stiffness cases. The remaining parameters are fixed as follows: division rate in the ventricular zone *G*_vz_ = 120, and stiffness ratio *β_μ_* = 8 for CS and 3 for VS.

### How does cell proliferation in the OSVZ affect the cell density and folding evolution?

After we have seen the effect of cell proliferation in the OSVZ on the final folding pattern, we investigate its effect on the evolution of both cell density and folding morphology. Figure 6 shows the temporal evolution of the maximum cell density in the domain and the folding evolution between time steps 260 and 380 for different initial division rates in the OSVZ *G*_osvz_ and a constant division rate in the VZ *G*_vz_ = 120. Again, we consider both cases constant and varying cortical stiffness. The folding evolution value quantifies the ratio between the outer perimeter at time step *t* and the initial perimeter as demonstrated in Figure 6A. Increasing the initial division rate in the OSVZ leads to a significant increase in both the cell density and folding evolution. Consequently, for the case of *G*_osvz_ = 30, the cell density reaches the highest value of 1100mm^-2^ corresponding to a folding evolution of 1.25. For the case of *G*_osvz_ = 0, in contrast, the cell density does not even exceed the migration threshold. The kink observable at time step 360 for the varying cortical stiffness case and an initial division rate in the OSVZ *G*_osvz_ = 20 indicates that the mechanical instability has occurred, i.e., the first gyri and sulci start to appear. Comparing the cases *G*_osvz_ = 20 and *G*_osvz_ = 30 shows that the instability occurs earlier with increasing initial division rate in the OSVZ. Thus, the ORGC proliferation decreases the time required to reach the final folding pattern. In general, the differences between the results for the constant and varying cortical stiffness cases are minor. However, the curves for the varying cortical stiffness case rise faster than for the constant case. We note that we could not generate results for the initial division rate in the OSVZ *G*_osvz_ = 30 and the constant cortical stiffness case due to numerical issues.

**Figure 6.**
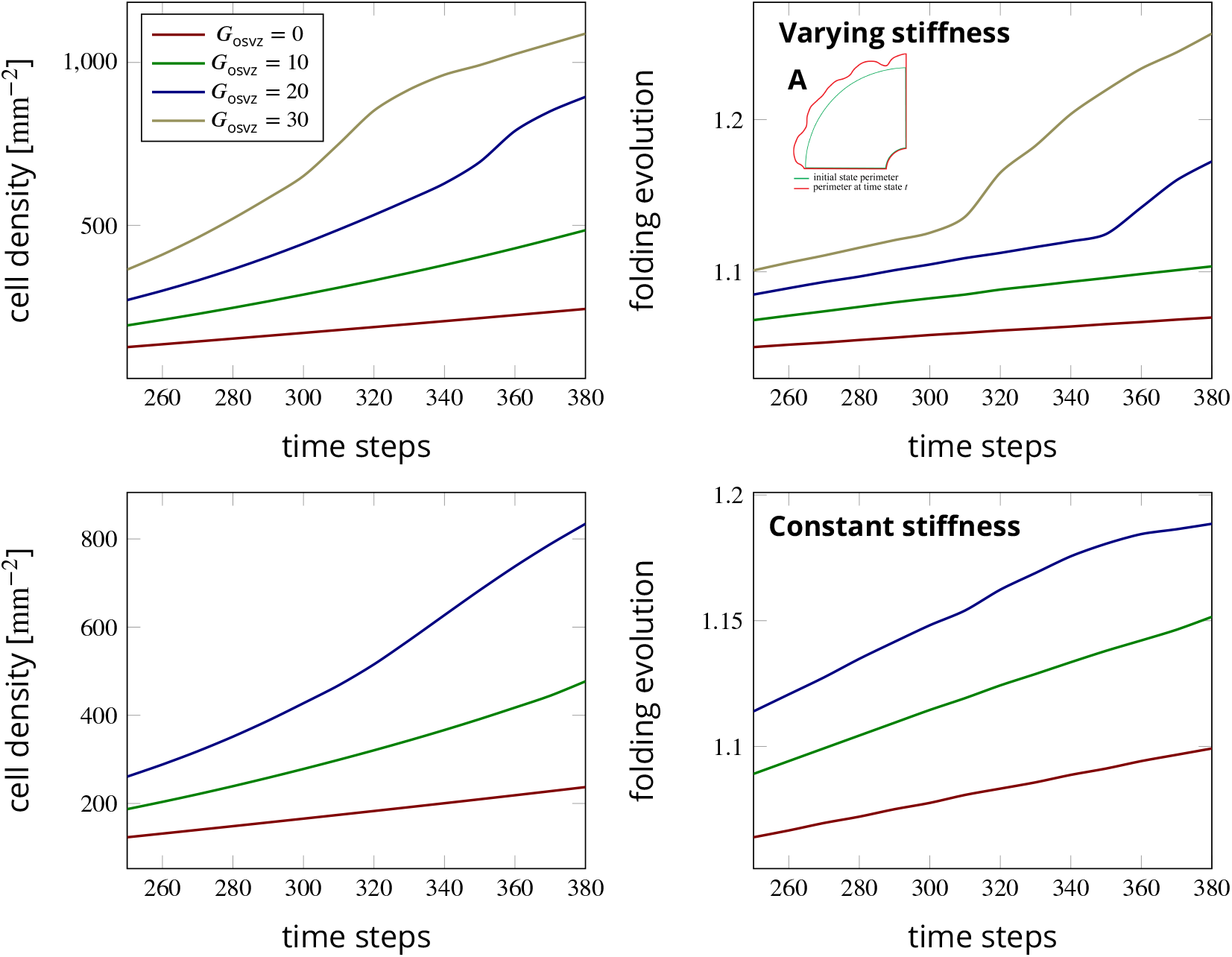
Temporal evolution of the maximum cell density and the folding evolution (the current outer perimeter divided by the initial perimeter as indicated in subfigure A) at a constant division rate in the ventricular zone (VZ) *G*_vz_ = 120 and different initial division rates in the outer subventricular zone (OSVZ) *G*_osvz_. The results in the top row correspond to the varying cortical stiffness case with a stiffness ratio of 3. The results in the bottom row correspond to the constant cortical stiffness case with a stiffness ratio of 8.

### Which proliferation zone is more influential for fetal human brain development?

To answer the question whether one of the proliferation zones (in the VZ or OSVZ) is more important for cellular brain development and cortical folding, we have implemented four sets of division rates for both the VZ and the OSVZ. The first set assumes a high division rate in the VZ *G*_vz_ and a low initial division rate in the OSVZ *G*_osvz_. For the following parameter sets, we gradually descrease *G*_vz_, while we increase *G*_osvz_. Figure 7 shows the corresponding results for the maximum cell density in the domain and the folding evolution between time steps 250 and 450 for both cases constant and varying cortical stiffness. Unexpectedly, the cell density for the set (*G*_vz_ = 30, *G*_asvz_ = 30) rises faster than the one for the set (*G*_vz_ = 120, *G*_osvz_ = 0). Indeed, not only the cell density is affected by increasing the division rate in the OSVZ (at the expense of decreasing it in the VZ), but also convolutions (cortical folds) appear earlier, as illustrated in the curve for the folding evolution (Figure 7, right). Our results thus indicate an unproportionally strong effect of ORGC proliferation in the OSVZ on cellular brain development and cortical folding. We attribute this observation to the the larger volume occupied by the OSVZ compared to the VZ. Concerning the difference between the results for the constant and varying cortical stiffness cases, the curves for a varying stiffness rise faster than for a constant stiffness.

**Figure 7.**
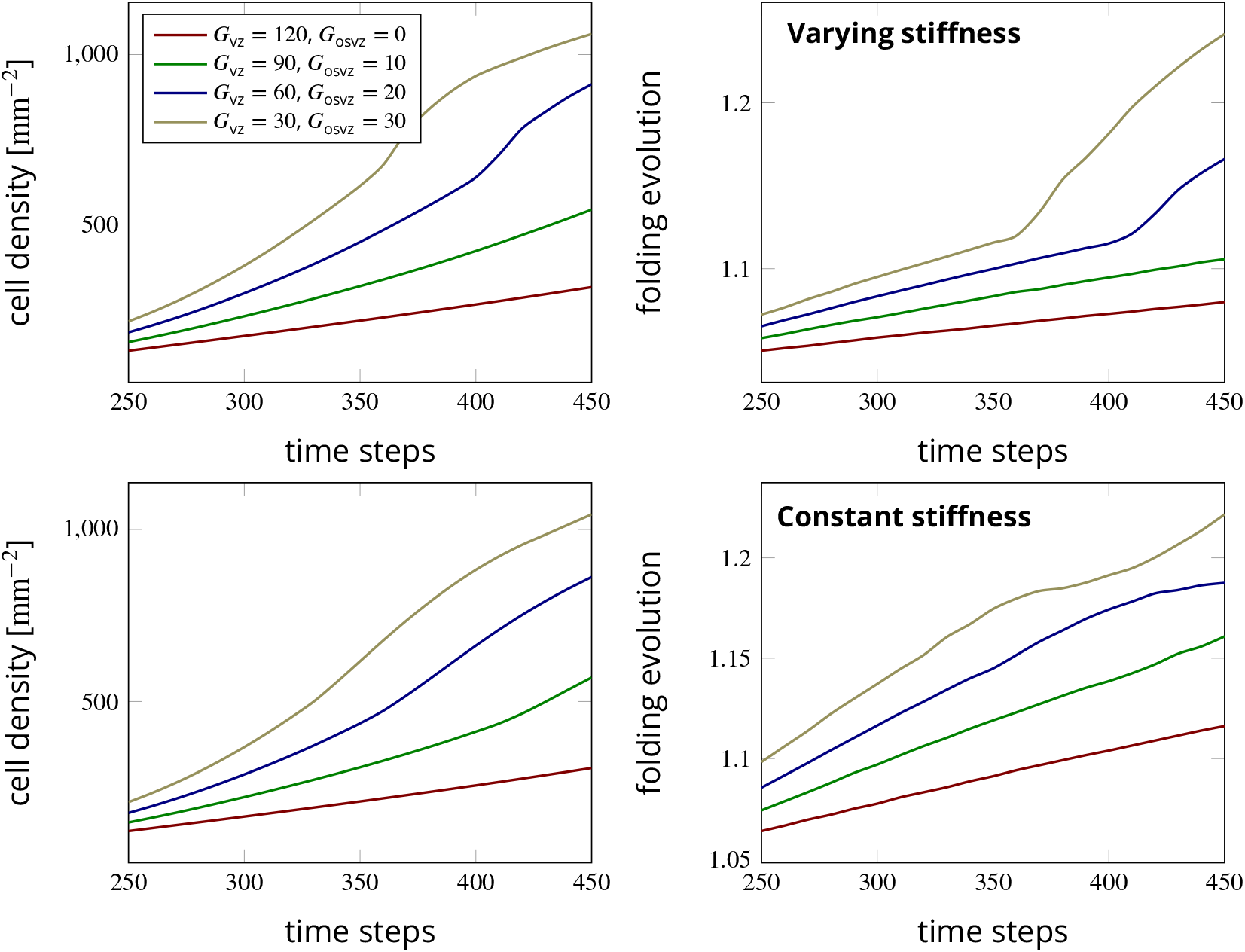
Temporal evolution of the maximum cell density and the folding evolution for different division rates in the ventricular zone *G*_vz_ and outer subventricular zone *G*_osvz_. The results in the top row correspond to the time-varying cortical stiffness case with a stiffness ratio of 3. The results in the bottom row correspond to the constant cortical stiffness case with a stiffness ratio of 8.

### How does the mitotic small translocation behavior of ORGCs affect brain development?

The final simulation parameter we would like to study is the MST factor that mimics the mitotic small translocation behavior of ORGCs. For this parameter study, we limit ourselves to the varying cortical stiffness case with a division rate in the VZ of *G*_vz_ = 30 and an initial division rate in the OSVZ of *G*_osvz_ = 30. Figure 8 shows the temporal evolution of the maximum cell density and the cortical folding evolution between time steps 250 and 430 for different values of the MST factor. Our simulations show that with increasing MST factor, the value of the maximum cell density in the domain increases exponentially. Since the MST factor incorporates the expansion of the OSVZ with time, these results are consistent with the observation in the previous section showing an increasing volume space of the OSVZ.

**Figure 8.**
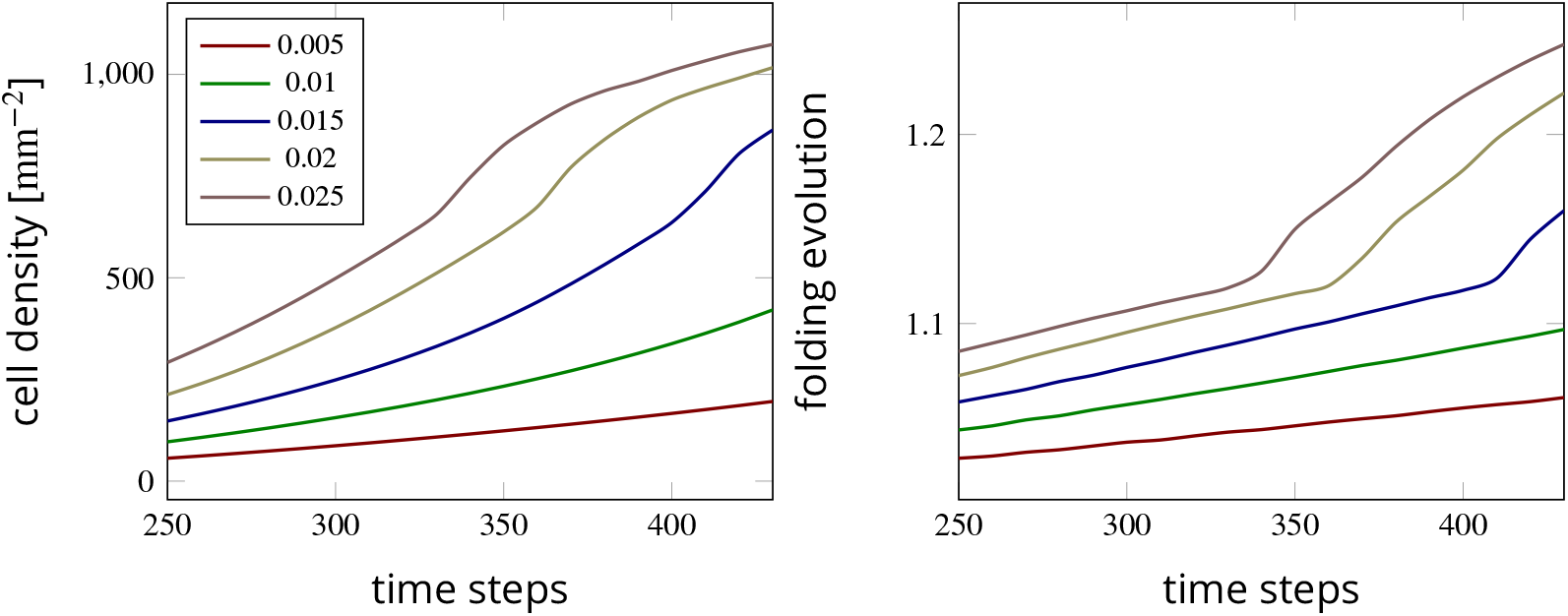
Temporal evolution of the maximum cell density and the folding evolution for different values of the mitotic small translocation (MST) factor. The results correspond to the varying cortical stiffness case with a stiffness ratio of 3, a division rate in the ventricular zone (VZ) of *G*_vz_ = 30, and an initial division rate in the outer subventricular zone (OSVZ) of *G*_osvz_ = 30.

### Is the OSVZ affected by cortical folding?

After we have discussed the effect of the OSVZ on the formation of cortical folds, we will now discuss the opposite – the effect of cortical folding on the OSVZ. In the intermediate stage of human brain development, before cortical folds emerge, the OSVZ has a constant thickness throughout the entire domain, as demonstrated in Figure 9, left, where the OSVZ appears in red color. However, according to our simulations, this quickly changes after the first folds start to emerge. The thicknessofthe OSVZ starts to vary and becomes thicker beneath gyri and thinner beneath sulci, as shown in Figure 9, right. This result is consistent with what was previously observed in the human brain ***Kostović et al. (2002)***. While it is to date not clear whether this phenomenon is rather the cause or the result of cortical folding, our study clearly supports the latter. From a mechanical perspective, it seems that the forces generated due to the underlying cellular mechanism, do not only fold the cortical layer but also lead to undulations in the deeper zones. Still, the deeper subcortical layers (i.e ISVZ and VZ) remain equally smooth as the ventricular surface.

**Figure 9.**
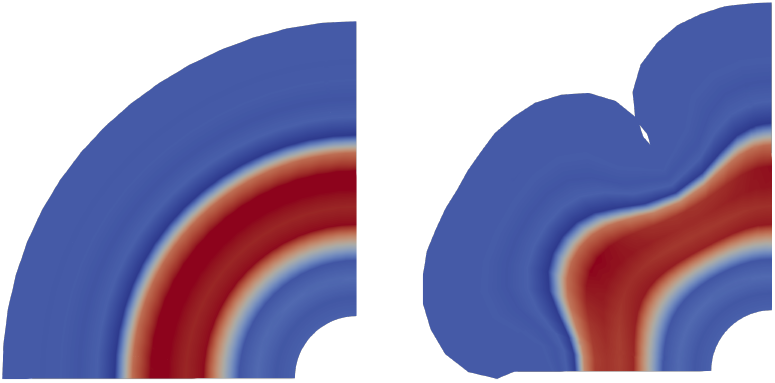
The effect of cortical folding on the OSVZ. While the OSVZ has a constant thickness before cortical folds emerge (left), it later becomes thicker beneath gyri than beneath sulci (right).

### Are cortical folds affected by regional proliferation variations in the OSVZ?

Previous studies have emphasized the existence of variations in the ORGC proliferation rate in different brain regions ***Hansen et al. (2010)***. Some of those have suggested that the proliferation rate is higher beneath gyri than beneath sulci ***Borrell (2018)***. To mimic this phenomenon, we apply a varying division rate by reformulating equation 8,

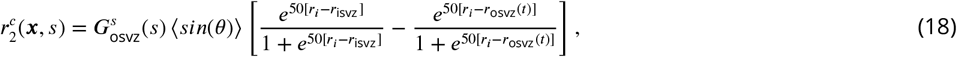

where 〈〉 are the Macaulay brackets, and ***θ*** is the angle between the x-axis and the point position vector. Figure 10 shows the development of cortical folds between time steps 325 and 450 for the varying cortical stiffness case, a division rate in the VZ *G*_vz_ = 120 and an initial division rate in the OSVZ of *G*_osvz_ = 20. The top row highlights the cell density distribution and the bottom row the locally varying division rate in the OSVZ. We observe that the cortex layer undergoes great surface expansion and the cortical folds indeed develop faster above the regions of higher division rates. Furthermore, the folds become more complex and have a higher cell density compared to regions nearby. However, the gyri and sulci are distributed equally between the different regions – regardless of the division rate. Still, sulci deepen faster above the regions with higher division rates. This simulation result is consistent with the previously found remarkable surface expansion above the regions with higher proliferation in the OSVZ ***Llinares-Benadero and Borrell (2019)***.

**Figure 10.**
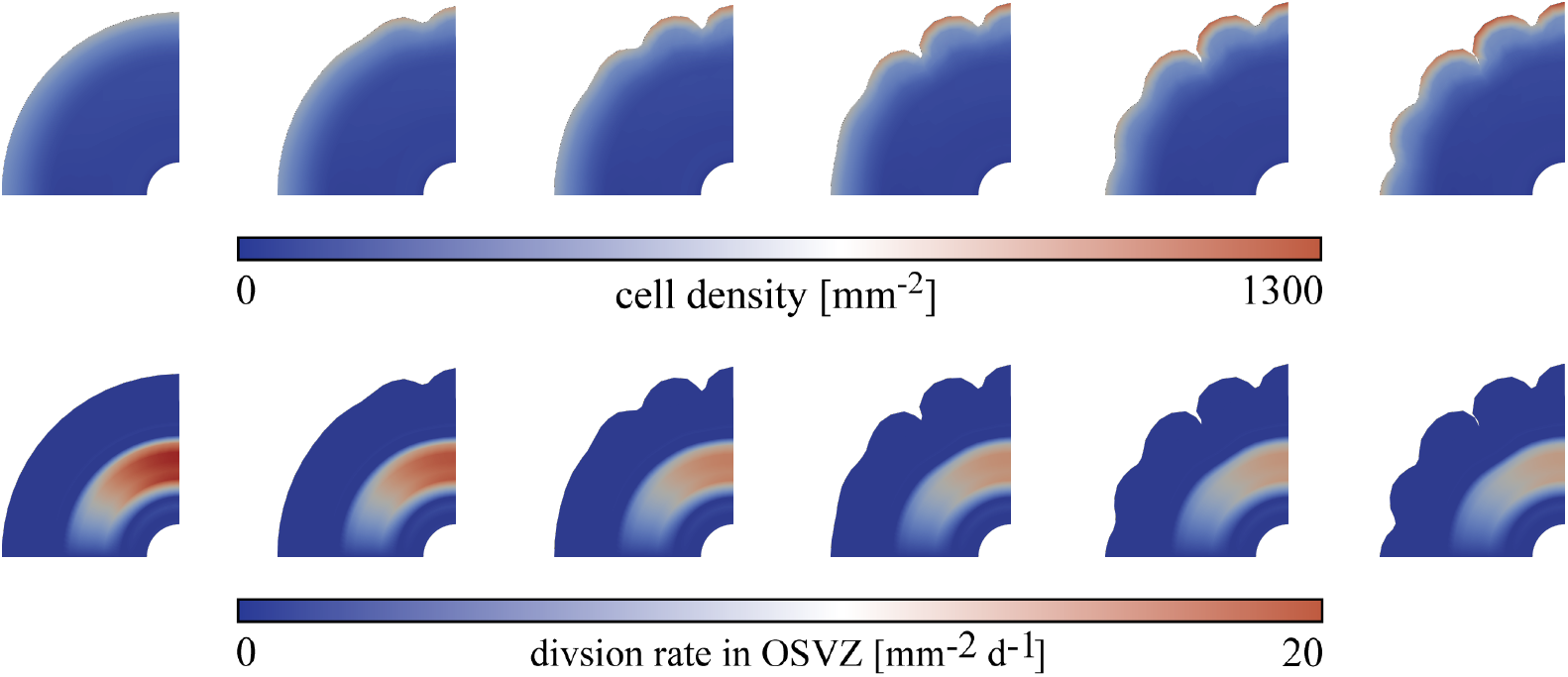
The development of cortical folds between time steps 325 and 450 for the varying cortical stiffness case and initial division rate in the VZ *G*_vz_ = 120 and varying division rates in the OSVZ with an initial value of 20. The top row shows the cell density distribution, and bottom row shows the actual division rate in the OSVZ.

## Conclusion

In this work, we have made use of a computational model for brain growth to provide insights into the role of the outer subventricular zone (OSVZ) – a unique additional proliferating zone in humans – during cortical folding in the developing brain. By using computational tools, we have addressed different open questions with the aim of valuably supplementing classical experimental approaches. Our simulations have systematically demonstrated how the proliferation in the OSVZ enhances the complexity of cortical folding patterns. An important decisive factor controlling the emerging folding pattern is the tissue stiffness ratio between the cortex and the subcortical plate. As the cortical stiffness appears to depend on the cell density, this is directly related to the cell pro-liferation in deeper zones. Our results show that the existence ofthe OSVZ particularly triggers the emergence of secondary mechanical instabilities leading to more complex folding patterns. Furthermore, the proliferation of outer radial glial cells (ORGCs) reduces the time required to induce the mechanical instability and thus cortical folding. Interestingly, our simulation results suggest that the generated mechanical forces not only ‘fold’ the cortex but also deeper subcortical zones including the OSVZ, which becomes thicker beneath gyri and thinner beneath sulci as a result of cortical folding. In turn, we did not find any relation between regionally varying ORGCs proliferation and the location of emerging sulci and gyri. Consequently, our physics-based analyses suggest that regional differences in the thickness of the OSVZ are rather a result of than a cause for cortical folding. Still, locally increased proliferation in the OSVZ leads to emergence of deeper sulci.

In conclusion, our physics-based computational modeling approach has proven valuable to predictively assess the links between cellular mechanisms during human brain development and cortical folding - the classical hallmark of the human cortex at the organ scale. It has allowed us to systematically assess the role of the OSVZ during human brain development and its effect on the cortical folding process. In the future, the computational framework can be used to not only better understand physiological brain development but also pathological processes - especially those involving abnormal cortical folding patterns. The computational model is able to shed new light on the interplay between the multiple processes at different scales and can help identify the main controlling parameters. However, it can only complement, not substitute, sophisticated experimental approaches that are still needed to answer questions, e.g., regarding the functional difference between progenitor cell types and corresponding lineage decisions.

## Acknowledgements

We would like to cordially thank Bettina Seydel for digitalizing the histological sections. In addition, we gratefully acknowledge the funding by the Deutsche Forschungsgemeinschaft (DFG, German Research Foundation) through the grant BU 3728/1-1 to SB.

## Author contributions statement

S.B. and M.S.Z. conceptualized the study and developed the model. S.B. acquired funding. M.S.Z. implemented the computational model, performed the simulations, and analyzed the results. I.B. provided the human fetal brain sections and contributed to data analysis. M.S.Z. visualized the results and wrote the first draft. S.B supported visualization and oversaw the writing process. All authors discussed the results, and reviewed and edited the manuscript.

## Data availability

The datasets and code generated and/or analyzed during the current study are available on GitHub: https://github.com/SaeedZarzor/brain-development.git

